# Virome and nrEVEome diversity of *Aedes albopictus* mosquitoes from La Reunion Island and China

**DOI:** 10.1101/2022.08.24.505081

**Authors:** Umberto Palatini, Niccolò Alfano, Rebeca Lejarazu Carballar, Xiaoguang Chen, Helene Delatte, Mariangela Bonizzoni

## Abstract

**Background:** *Aedes albopictus* is a public health threat for its worldwide spread and ability to transmit arboviruses. Understanding mechanisms of mosquito immunity can provide new tools to control arbovirus spread. The genomes of *Aedes* mosquitoes contain hundreads of nonretroviral endogenous viral elements (nrEVEs), which are enriched in piRNA clusters and produce piRNAs, with the potential to target cognate viruses. Recently, one nrEVE was shown to limit cognate viral infection through nrEVE-derived piRNAs. These findings suggest that nrEVEs constitute an archive of past viral infection and that the landscape of viral integrations may be variable across populations depending on their viral exposure.

**Methods:** We used bioinformatics and molecular approaches to identify known and novel (i.e. absent in the reference genome) viral integrations in the genome of wild collected *Aedes albopictus* mosquitoes and characterize their virome.

**Results:** We showed that the landscape of viral integrations is dynamic with seven novel viral integrations being characterised, but does not correlate with the virome, which includes both viral species known and unknown to infect mosquitoes. However, the small RNA coverage profile of nrEVEs and the viral genomic contigs we identified confimed an interaction among these elements and the piRNA and siRNA pathways in mosquitoes.

**Conclusions:** Mosquitoes nrEVEs have been recently descrived as a new form of heritable, sequence-specific mechanism of antiviral immunity. Our results contribute to understanding the dynamic distribution of nrEVEs in the genomes of wild *Ae. albopictus* and their interaction with mosquito viruses.

---

Following its rapid global spread, the Asian tiger mosquito *Aedes albopictus* has emerged as a serious public health threat for its ability to transmit over twenty human pathogenic viruses, including dengue, chikungunya and Zika viruses, and the nematode *Dirofilaria immitis* (1, 2). Besides arboviruses, mosquitoes can be infected with viruses that are not able to replicate in vertebrate cells and are thus called insect-specific viruses (ISVs) (3, 4). ISVs are phylogenetically related to arboviruses, for instance, the *Flaviviridae* family contains two ISVs groups called classical Insect-Specific Flaviviruses (cISFVs) and dual-host ISFVs, respectively. Dual-host ISFVs are phylogenetically nested within mosquito-borne viruses, whereas cISFs are a monophyletic group thought to be an ancestral lineage of Flaviviruses (5). ISVs are also abundant in the *Rhabdoviridae, Bunyaviridae* and *Reoviridae* families and are present, but have not been so frequently identified yet, in the *Togaviridae* family (6). Phylogenetic analyses within the *Bunyaviridae* family support the hypothesis that arboviruses of this viral family evolved from ISVs by overcoming the barriers preventing infection of vertebrates. ISVs have also been proposed as new biological control agents against arboviruses based on the observation that some ISVs, such as Eilat Virus in *Ae. albopictus*, alter mosquito vector competence by upregulating the antiviral immune responses or through superinfection exclusion (7). Despite the recognized significance of ISVs, knowledge on the virome diversity of wild-collected *Ae. albopictus* mosquitoes is still limited (8–11). *Aedes albopictus* populations are usually classified into native, old and invasive (1). Native populations are endemic in the species home range in Southeast Asia; old populations can be found in the islands of the Indian Ocean, which were colonized in the ‘800 century, and invasive are populations resulting from overlapping and chaotic incursions which took place in both tropical and temperate regions of the world in the past 50-60 years (12).

Both ISVs and arboviral infections elicit RNA interference (RNAi) pathways in mosquitoes. RNAi depends on the production of small (s)RNAs, which are classified on the basis of their size, biogenesis and interacting proteins into microRNAs of ∼20– 22 bp; small interfering (si)RNAs of ∼20– 22 bp,, and PIWI interacting (pi)RNAs of ∼25-30 bp (13). miRNAs derive from hairpin-like structure forming in long single-stranded RNA molecule, siRNAs are processed from long double stranded RNAs, while piRNAs are produced both in the nucleus (primary piRNAs) from *ad hoc* genomic regions called piRNA clusters and in the cytoplasm (secondary piRNAs) from self-amplification of primary piRNAs after target recognition and processing (14). While during viral infection of mosquitoes, siRNAs are consistently found and homogeneously cover infecting viral genomes, production of virus-derived miRNAs is controversial and the profile of piRNAs appears to be tissue-specific and both viral and host-specific (15–17). On this basis, the pattern of small RNAs, *in primis* siRNAs, has been used to reconstruct the sequences of viruses infecting mosquitoes and other animals (16, 18, 19).

piRNA clusters of *Aedes* mosquitoes, but not those of *Drosophila melanogaster* nor Anophelinae, are enriched in non-retroviral RNA virus endogenous elements (nrEVEs) (20–24). The genome of *Ae. albopictus* hosts a total of 456 nrEVEs (20). The most represented viral families from which nrEVEs originate are *Rhabdoviridae* (Rhabdo-nrEVEs), *Flaviviridae* (Flavi-nrEVEs) and the newly discovered *Chuviridae* (20, 23, 25), besides unclassified viruses identified in arthropods but whose host range is still unknown. *Aedes albopictus* nrEVEs produce piRNAs which are antisense with respect to cognate viruses suggesting antiviral activity (21). Reduction of cognate viral infection due to nrEVE-derived piRNAs has been shown for selected nrEVEs of both *Ae. aegypti* and Aag2 cells (6, 26, 27). Despite an antiviral activity of nrEVEs have not been proven in *Ae. albopictus*, results in *Ae. aegypti* suggest that nrEVEs are involved in shaping the relationships among mosquitoes, ISVs and, possibly, arboviruses. However, information on the co-occurrence between nrEVEs and ISVs in geographic mosquito populations is lacking. To address this knowledge gap, we sampled *Ae. albopictus* mosquitoes from the native home range in Southern China and in La Reunion Island (1) and combined whole genome (WG) and small RNA sequencing approaches to characterize the concomitant patterns of viral integrations and their the small RNA profile while also detecting the mosquito virome.

## 2. Materials and Methods

### 2.1. Wild samples collection and sequencing

Adult *Ae. albopictus* individuals were collected in public areas from four sites in La Reunion Island (France) and two sites in the Guangzhou prefecture (China), which did not require ethical approval (**Fig. 1**). Mosquitoes collected in La Reunion Island were immediately processed in pools of 10 females and homogenized in DNA/RNA-shield (Zymo Research, Irvine CA, USA) with a Kimble electric pestle (DWK Life Sciences, Mainz, Germany). Samples were stored at 4°C before being transferred to the University of Pavia where each pool was divided into two volumes that were used for either DNA or RNA extraction. DNA was extracted from mosquito pools using the DNeasy Blood and Tissue Kit (Qiagen, Hilden, Germany), according to manufacturer’s protocol and stored at -20 °C. RNA was extracted using a standard protocol based on Trizol (Ambion/Invitrogen, Waltham MA, USA), treated with DNase I (Sigma-Aldrich, St. Louis MO, USA) and stored at -80 °C. Nucleic acids from three pools were further pooled to generate a sample. For each locality we processed two samples, which were sent to Biodiversa (Rovereto, Italy) for library preparation and sequencing resulting in the analyses of a total of 60 mosquitoes per site (**Table S1**). WG and Small RNA libraries were prepared using the Nextera DNA Library Preparation Kit for paired end reads (Illumina, San Diego CA, USA) and the NEBNext Small RNA Library Prep Set (New England Biolabs, Ipswich MA, USA), respectively. DNA and small-RNA Libraries were sequenced on an Illumina HiSeq 2500 instrument.

**Fig. 1.**
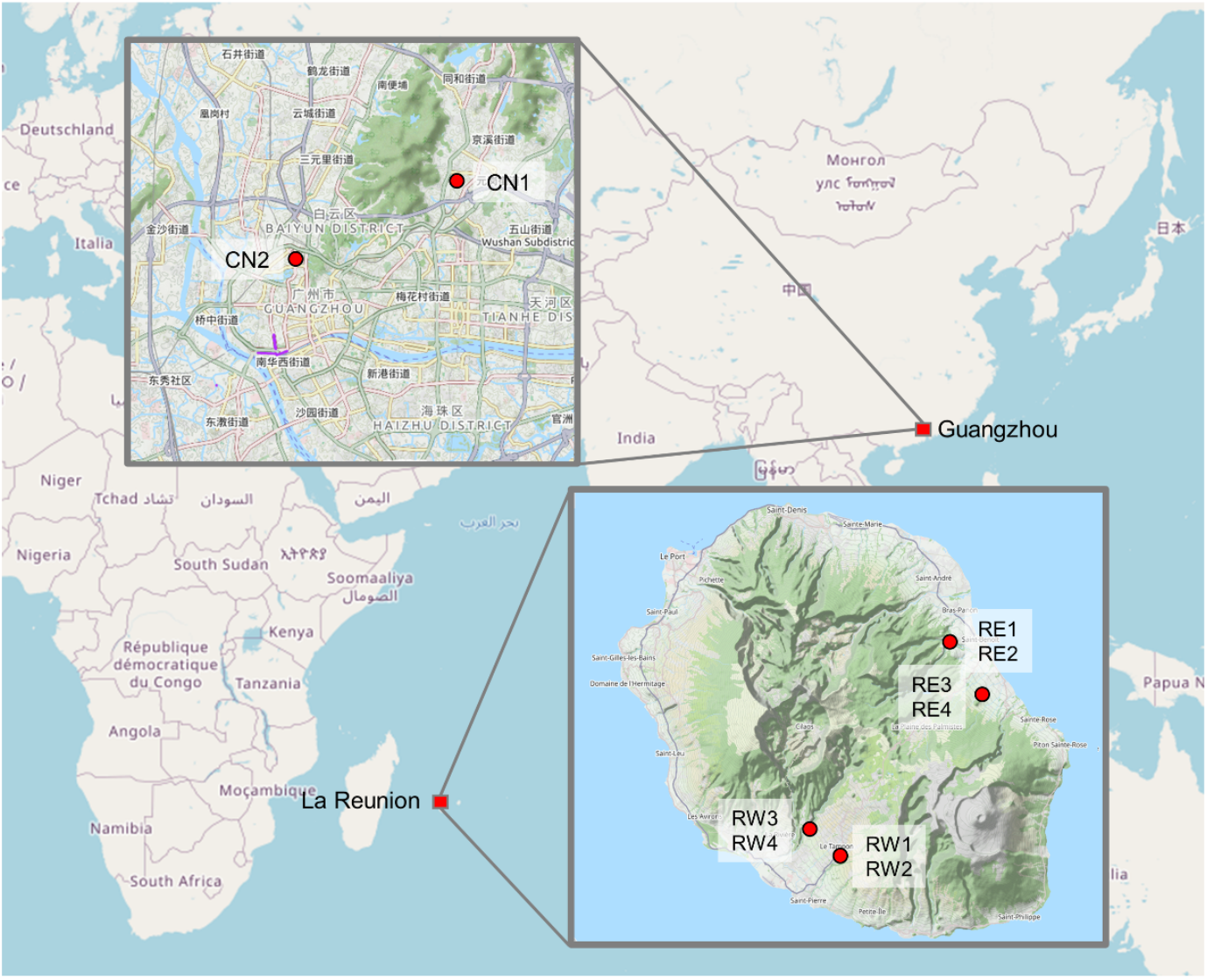
Collection sites of mosquitoes in Guangzhou (China) and La Reunion (France). In Southern China, two localities, CN1 and CN2, were sampled. In La Reunion, we sampled in four localities, two on the East part of the island (RE) and two on the West (RW). For each site two samples, each of 30 mosquitoes, were analyzed.

Samples from China were also processed in pools of 10 mosquitoes each. DNA was extracted using the Universal Genomic DNA extraction kit (Takara, Kusatsu, Japan) according to the supplier protocol and stored at -20 °C. RNA was extracted using a custom protocol based on Trizol (Ambion/Invitrogen, Waltham MA, USA), treated with DNase I (Takara, Kusatsu, Japan) and stored at -80 °C. DNA and RNA. As for samples from La Reunion, 3 pools of the nucleic acids of 10 mosquitoes each were further pooled and sent to the Beijing Genomics Institute (BGI), China, for library preparation and sequencing (**Table S1**). DNA WGS libraries were prepared with the Nextera DNA Library Preparation Kit (Illumina, San Diego CA, USA) and were sequenced on an Illumina HiSeq X instrument. Small RNA libraries were prepared with the TruSeq Small RNA Library Prep Kit (Illumina, San Diego CA, USA) and sequenced on an Illumina HiSeq 4500 sequencer.

### 2.2. nrEVEome

WGS data were analyzed to identify the landscape of nrEVEs of wild-collected mosquitoes using as reference the nrEVEs that we previously annotated in the *Ae. albopictus* reference genome (AalbF2) (20). Reads were checked and cleaned with with FastQC v. 0.11.9 (https://www.bioinformatics.babraham.ac.uk/projects/fastqc/) and trimmomatic v. 0.40 (28), respectively, and mapped on the AalbF2 genome using the SVD pipeline as previously described. Given that we sequenced pools of 30 individuals, we estimated the frequency of nrEVEs in each pool based on read coverage. Briefly, single copy orthologs were identified in the AalbF2 genome assembly using BUSCO v5.3.1 (29), configured with the diptera_odb10 database. Normalized depth values of a selected set of 359 orthologs, which satisfied a minimum BUSCO score of 1000 (**table S2**), and of all the reference nrEVEs were extracted using FeatureCounts in the SubRead package (30). Average normalized depth for single copy orthologs was calculated for each sample and used as a reference to estimate nrEVE allelic frequency in pooled samples (**Fig. S1**). nrEVEs were considered “present” in a sample when the estimated allelic frequency was higher than 0.2. The Principal Component Analysis (PCA) classifying our samples based on their pattern of nrEVEs was computed using the Exponential family PCA method in the LogisticPCA R (31) package and plotted with ggplot2.

### 2.3. Characterization of novel viral integration sites in wild collected mosquitoes

The Vy-PER and ViR pipelines were run as previously described (32) to identify in the DNA of wild-collected mosquitoes nrEVEs which are absent in the AalbF2 assembly. Bioinformatic predictions of each of the newly identified viral integrations were molecularly confirmed by PCR using the DreamTaq Green PCR Master Mix 2X (Thermo Fisher) and nrEVE specific primers (**Table S3)**. PCR products were cloned into *E. Coli* OneShot TOP10 chemically competent cells (Invitrogen) using the TOPO TA Cloning Kit for Sequencing (Thermo Fisher, USA). Plasmids containing the PCR product were extracted with the QIAprep Spin Miniprep Kit (Qiagen, Germany). Plasmid products were Sanger sequenced by Macrogen Europe (Amsterdam, The Netherlands) using the M13F and M13R plasmid primers.

The presence of the newly identified nrEVEs was also tested by PCR on single mosquitoes of laboratory strains derived from mosquitoes sampled in Canton (China), Chiang Mai (Thailand), La Reunion Island, Crema (Italy) and Tapachula (Mexico). DNA was extracted from single mosquitoes using the Wizard genomic DNA purification kit (Promega) and PCRs were performed as described above

### 2.4. Virome analyses

Raw small-RNA (sRNA) sequencing data were checked for quality with FastQC v. 0.11.9 (https://www.bioinformatics.babraham.ac.uk/projects/fastqc/); the presence of adapters was confirmed with DNApi v. 1.1 (33). Adapters and bases with baseQ inferior to 20 were trimmed from the reads using Cutadapt v. 2.9 (34), also removing clean reads smaller than 10 bp and longer than 50 bp. Low complexity reads were excluded with the Duster.pl script from the NGS-toolbox package (https://sourceforge.net/p/ngs-toolbox/wiki/Home/). Briefly, reads were mapped to the *Ae. Albopictus* genome (AalbF2 assembly) after masking nrEVEs using BEDtools maskfasta v. 2.28 (35). Small-RNA reads were mapped on the masked genome with Bowtie v. 1.1.2 (36) optimizing the parameters for small-RNA reads (-n 1 -l 18 --best). The reads that did not map to the *Ae. albopictus* genome were extracted with SAMtools view v. 1.4 (37) and converted to fastq with SAMtools fastq. Contigs were reconstructed from unmapped sRNA reads using the Oases pipeline (38) with Velvet v.1.2.10 (https://www.ebi.ac.uk/~zerbino/velvet/) testing k-mer lengths from 13 to 31 bp. Sequences assembled by Oases and longer than 100bp were filtered to remove nrEVEs-derived assemblies using BLASTn v. 2.6.0 (39) by comparing all the assembled sequences against nrEVEs annotated in AalbF2 (20). Contigs with a percentage of identity higher than 90% and covering at least 50% of an existing nrEVE were considered nrEVEs assemblies rather than viral contigs. Redundant assemblies were clustered using CD-HIT v. 4.8.0 (40) with the following options: -c 0.9 -n 8 -d 60 -g 1. The clustered assemblies were screened against the entire NCBI nr peptides database (https://www.ncbi.nlm.nih.gov/refseq/about/nonredundantproteins/, downloaded in march 2020) using DIAMOND v. 0.9.31 (41) BLASTx algorithm with an e-value cutoff of 1e^-5^. DIAMOND hits were loaded on MEGAN6 community edition v. 6.19.9 (42). MEGAN6 was run with the naive LCA algorithm. Assemblies assigned to viruses by MEGAN6 were manually validated against the nr database using BLASTx (39).

### 2.5. small RNA profile of viral contigs and novel nrEVEs

Selected assembled viral contigs, including viral contigs of at least 1500bp and Flavivirus-like viral contigs shorter than 500 bp, were used for the realignment of sRNA reads from the samples in which the contigs were detected. Clean sRNA reads were mapped using Bowtie v. 1.1.2 (36) as described above. Sequences between 20–22 bp and 25–30 bp were filtered as siRNAs/miRNAs and piRNAs, respectively, using BBMap reformat (sourceforge.net/projects/bbmap/). Small RNAs that mapped to viral contigs (hereafter called viral sRNA) were then re-mapped to the AalbF2 genome assembly, then split by orientation with samtools view and counted with BBMap pileup.sh (sourceforge.net/projects/bbmap/). The reads were categorized as either mapping exclusively on viral contigs or mapping on both viral contigs and nrEVEs. Reads size distributions were plotted on Python 3 using the Plotly v. 5.7.0 graphing library (43).

To understand the siRNA coverage of the novel viral integrations identified in wild-collected mosquitoes, the sRNA reads from each mosquito pool in which the viral integration had been identified were re-mapped to the viral integration using the methodology described above. Coverage profile of mapped piRNA and siRNA was plotted with GraphPad Prism 8 (www.graphpad.com).

## RESULTS

Adult *Ae. albopictus* mosquitoes were collected from four localities from La Reunion Island and two sites in the Guangzhou prefecture (China) (**Fig. 1**). Sampling sites included sites from the Eastern (Bras-Panon [RE1, RE2] and Saint-Rose [RE3, RE4]) and the western (Tampon [RW1, RW2] and Saint Pierre [RW3, RW4]) part of La Reunion Island, which experience low and high incidence of arboviral disease transmission, respectively (44). In the Guangzhou prefecture, we sampled mosquitoes from the Baiyun district (CN1 sample), which is located at the northern outskirts of prefecture and is a forested area known for its natural attractions and from the campus of the Southern Medical University (CN2 sample), an urban and densely populated area of the Guangzhou city, which is subjected to intense vector control interventions (45). From all sites, WGS and small RNA data were concomitantly produced from pools of 30 mosquitoes each (**Table S1**). WGS data were used to characterize the patterns of nrEVEs and test for the presence of new viral integrations. sRNA data were used to estimate the sRNA profile nrEVEs, including newly identified ones, and assemble viral contigs.

### 3.1 Patterns of nrEVEs across samples

Out of the 456 nrEVEs annotated in the *Ae. albopictus* genome, 222 nrEVEs appear to be absent in all samples, while 234 nrEVEs were estimated to be present based on read coverage with respect to mosquito single-copy ortholog genes. The majority of nrEVEs, which were not detected in any samples, included Chuviridae-like nrEVEs and nrEVEs from unclassified viruses followed by Rhabdovirus-like nrEVE. To determine if nrEVE patterns differ across samples based on their geographic origin, we used Principal Component Analysis (PCA). While results showed a clear separation between samples from China and La Reunion Island (**Fig. 2A**), within La Reunion, samples from the East and West coast did not group separately, consistent with high gene flow among populations of the island (44).

**Fig. 2.**
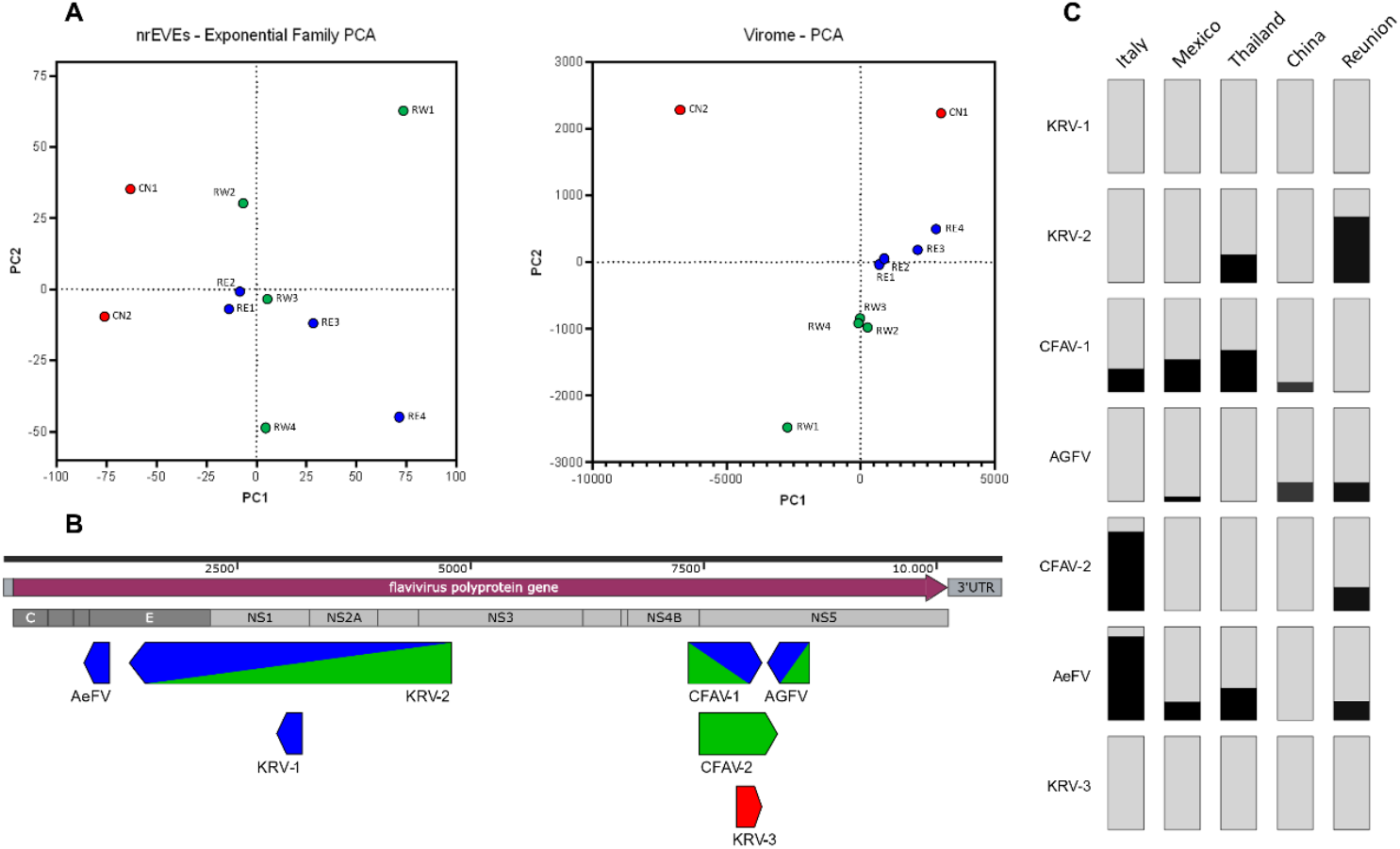
nrEVEome and virome of wild *Ae. albopictus* mosquitoes from China and La Reunion Island. (A) Principal Component Analysis based on the presence/absence of nrEVEs across samples or assembled viral contigs on the left and right panels, respectively. (B) Schemes of newly identified viral integrations with respect to the genome of a representative *Flavivirus*. Samples or new viral integrations from China are in red. Samples or new viral integrations from samples of the East and West side of La Reunion are in blue and green, respectively. (C) Bar plots showing the frequency of each newly identified viral integration in mosquitoes from Crema (Italy), Tapachula (Mexico), Chiang-Mai (Thailand) and Canton (China).

We further used the Vy-PER and ViR pipelines to align WGS data to the *Ae. albopictus* genome (AalbF2 assembly), identify unmapped reads containing viral sequences and characterize novel integration sites. A total of 7 new integrations were identified: 6 were detected in samples from La Reunion Island and one in Chinese mosquitoes (**Table 1, Fig. 2B, Table S1**). All newly identified nrEVEs derived from ISVs of the *Flaviviridae* family, including Kamiti River Virus (KRV), Cell Fusing Agent Virus (CFAV) and Aedes Flavivirus, with a percentage of nucleotide identity ranging from 67 to 78%. Additionally, these nrEVEs, which range in size between 814 and 4705 bp, encompass one or more viral proteins without breaks or rearrangement with respect to the original viral genome. Each of the newly identified nrEVE was molecularly validated by PCR using specific primers followed by Sanger sequencing (**Table S3**). Additional wild-caught mosquitoes from La Reunion Island and China, besides samples from Chiang Mai (Thailand), Crema (Italy) and Tapachula (Mexico) were used to assess the frequency of the newly identified nrEVE (**Fig. 2C**). Apart from KRV-1 and KRV-3 which were detected only in RE2 and CN1, respectively, all other newly identified nrEVEs were found in mosquitoes from both native and invasive populations, at frequency between 4% and 90% (**Fig. 2C**).

**Table 1:** List of novel viral integrations identified in wild-collected mosquitoes

| nrEVE ID | Virus | Contig length | Viral bp | % AA ID | Detected in |
| --- | --- | --- | --- | --- | --- |
| **KRV-1** | Kamiti River Virus | 814 | 309 | 74 | RE4 |
| **KRV-2** | Kamiti River Virus | 4705 | 3974 | 67 | RE4, RW2 |
| **CFAV-1** | Cell Fusing Agent Virus | 816 | 307 | 77 | RE2, RE3, RW2, CN1 |
| **AGFV** | Aedes Galloisi Flavivirus | 977 | 549 | 73 | RE3, RE4, RW2, RW4 |
| **CFAV-2** | Cell Fusing Agent Virus | 1623 | 891 | 71 | RW2, RW4 |
| **AeFV** | Aedes Flavivirus | 1046 | 321 | 77 | RE4 |
| **KRV-3** | Kamiti River Virus | 1882 | 947 | 74 | CN1 |

Since it is known that viral integrations are enriched in piRNA clusters and may produce piRNAs, we checked if, and where, piRNAs and siRNAs mapped on the newly detected nrEVE. Small RNAs in the size of piRNA, but not siRNAs were seen mapping across the novel viral integrations; coverage was not homogenous across the whole viral integration, but the overall profile was similar across samples (**Fig. 3**).

**Fig. 3.**
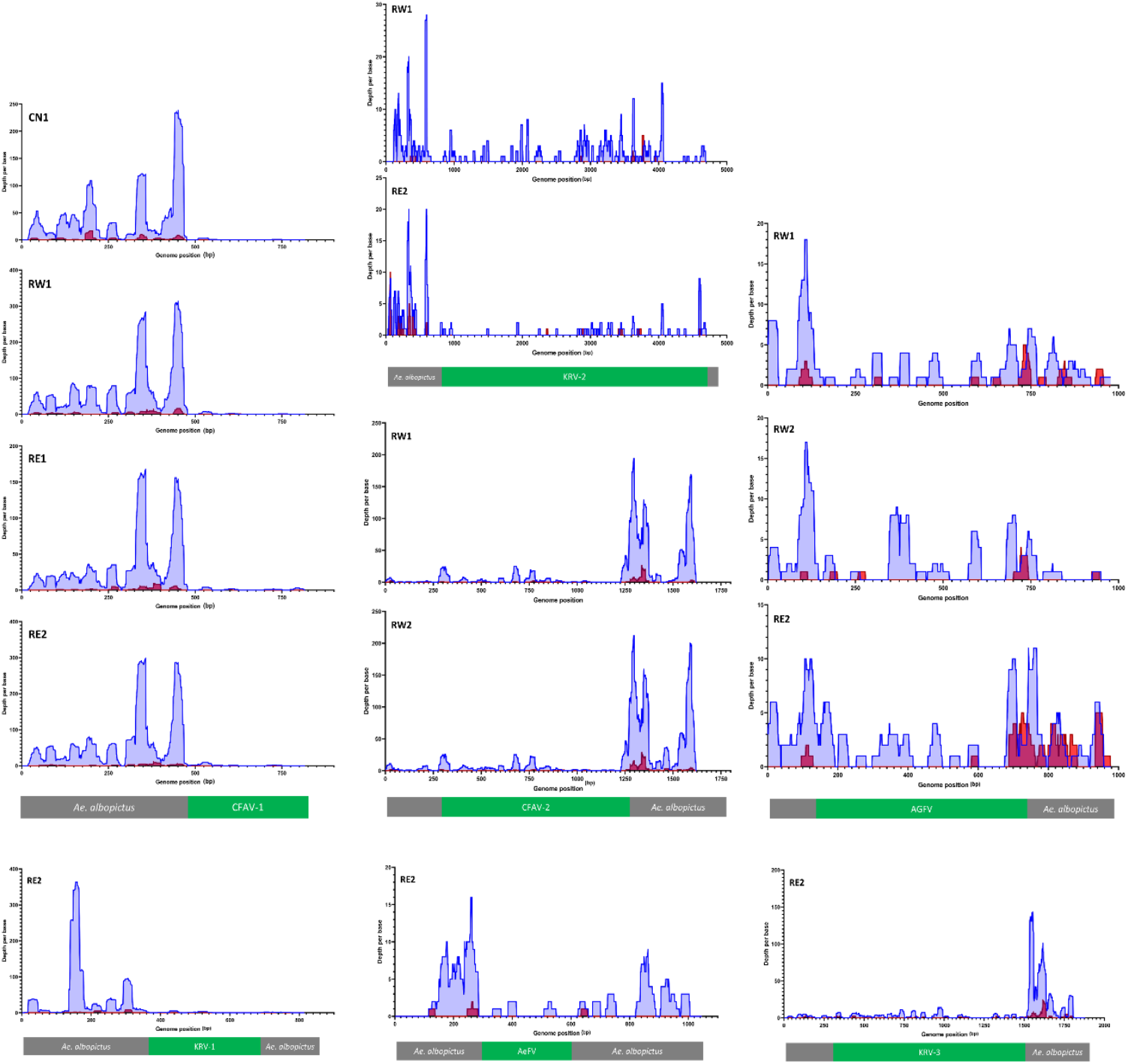
Small RNA coverage profile of newly identified nrEVEs. Coverage was calculated for siRNA/miRNAs (20-22nt) and piRNAs (26-30nt) and plotted on the DNA sequence of newly identified nrEVEs. Read peaks represent siRNA/miRNA coverage, while blue peaks piRNA coverage. Abbreviation on the left inside of each panel indicate the samples in which the nrEVEs was annotated and whose smallRNA were used to assess the profile.

### 3.2 The virome of Ae. albopictus mosquitoes

A total of 122 viral contigs corresponding to 25 viral species were assembled from small RNAs of Chinese mosquitoes. A total of 126 viral contigs corresponding to 26 viral species were assembled from small RNAseq data of samples from La Reunion (**Fig. 4**). Viral contigs ranged in size between 101 and 5847 bp and had a highly variable percentage of nucleotide identity (between 33 and 100%, with a median of 70%) with respect to known circulating viruses (**Fig. 4**). Most assembled contigs had similarities to *bona fide* arthropod viruses and included ISVs of the *Flaviviridae* family (i.e. CFAV, KRV, La Tina virus, and Aedes Flavivirus [AeFV]) (5, 7, 46), the *Mononegavirales* order (*Culex tritaeniorhynchus rhabdovirus* [CTRV] (47) and Merida virus (48) and Aedes Anphenvirus (49) from the *Rhabdoviridae* and *Xinmonoviridae* families, respectively); Yongsang bunyavirus 1, previously identified in *Aedes vexans* mosquitoes from Korea (50), Phasi-Charoen Like Virus (family *Phenuiviridae*), which is highly prevalent in both wild-caught *Ae. aegypti* and *Ae. albopictus* mosquitoes (51), along with less known viruses, which have been recently characterized from insect samples, such as Wuhan Mosquito Virus, Wenzhou sobemo-like virus, Guadalupe mosquito virus, Hubei mosquito virus, Chuvirus Mos8Chu0, Croada virus and Kaiowa virus (11, 52, 53). We did not assemble any contigs from arboviruses, except for a contig of 168 bp with sequence similarity to a member of the Vesicular stomatitis virus serogroup of the *Rhabdoviridae* family, the Yug Bogdanovac virus (YBV), which was initially isolated from sandflies (54), and we assembled from small RNAs of the RE1 sample from the Eastern side of La Reunion.

**Fig. 4.**
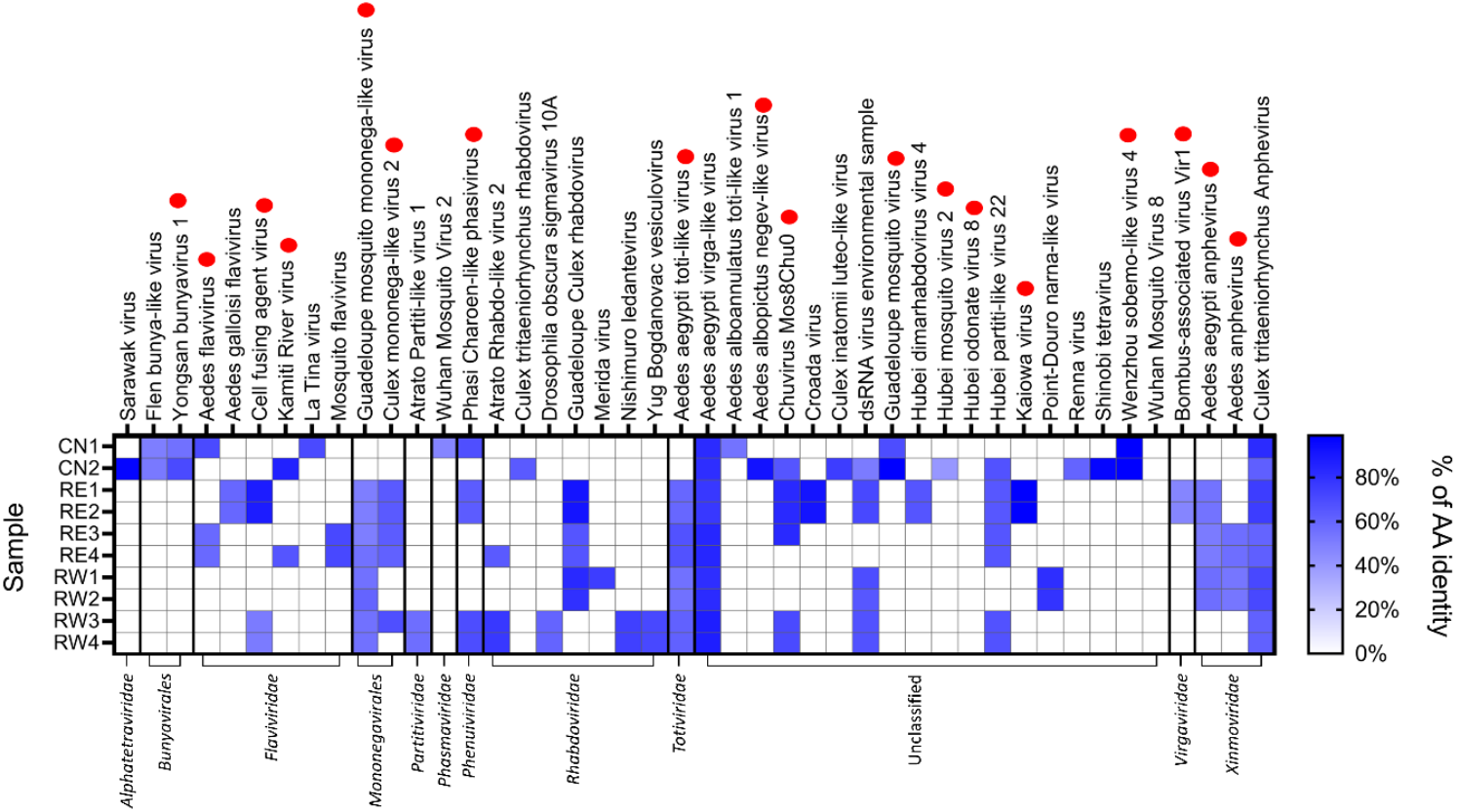
Diversity of viral contigs assembled from small RNAs of *Ae. albopictus* mosquitoes. Heat map showing the aminoacidic identity of assembled viral contigs with respect to cognate viruses. Viral contigs are shown in samples in which they were identified. Red dots indicate viral species which have been previously detected in wild-collected *Aedes* mosquitoes (9).

Because viral replication elicits RNAi and results in a small RNAs profile skewed towards siRNAs that target both strands of a viral genome, we checked this size signature on the viral species for which we assembled contigs of at least 1500bp, including Sarawak virus; Aedes aegypti toti-like virus; Aedes aegypti virga-like virus; Guadeloupe mosquito mononega-like virus,; Chuvirus Mos8Chu0, Point-Douro narna-like virus, Aedes albopictus negev-like virus and Shinobi tetravirus with a contig of 3450 bp (**Fig. 5**). Importantly, none of the assembled viral contigs matched to *Ae. albopictus* nrEVEs or newly identified nrEVEs. However, because endogenous piRNAs can trigger production of trailer and responder piRNAs from viral RNAs (55), we realigned piRNAs from the viral contigs to nrEVEs and tagged them as possible nrEVEs-derived small RNAs. A profile indicative of active viral infection was observed for the Sarawak, Shinobi tetravirus, the Aedes albopictus negev-like and, possibly, the Guadalupe mosquito mononega-like virus (**Fig. 5A**). On the contrary the smallRNA profile of the Aedes aegypti toti-like virus was skewed towards piRNAs, some of which may also derive from viral integrations despite the absence of sequence complementarity between the viral contig and nrEVEs (55). Furthermore, the profiles of contigs classified as Chuvirus Mos8Chu0, Aedes aegypti toti-like virus and Aedes aegypti virga-like virus were skewed to small RNAs of exclusively one strand and included smallRNAs which also map to nrEVEs. Because piRNAs against a single orientation of the viral genome with no siRNAs is a classic signature of nrEVEs, whereas siRNAs of positive and negative strand polarity are typical for viral DNA (vDNA) fragments, which have been shown to be formed after mosquito infection by a wide range of RNA viruses, possibly from their defective viral genomes (56–59), we cannot exclude that the smallRNA pattern observed for the Aedes aegypti toti-like virus contig represent an undetected novel nrEVE, whereas the profiles of the Chuvirus Mos8Chu0 and the Aedes aegypti virga-like virus are difficult to interpret. We also verified the profile of assembled flaviviral contigs shorter than 500 bp, which may represent vDNA fragments of flaviviruses or viral integrations which we were not able to identify from WGS data. The observed profiles with a prevalence of small RNAs in the size of piRNAs of only one orientation (**Fig. 5B**) points to nrEVEs, which are probably rare in the pool samples analyzed thus resulting in DNA reads below the threshold for novel nrEVE detection.

**Fig. 5.**
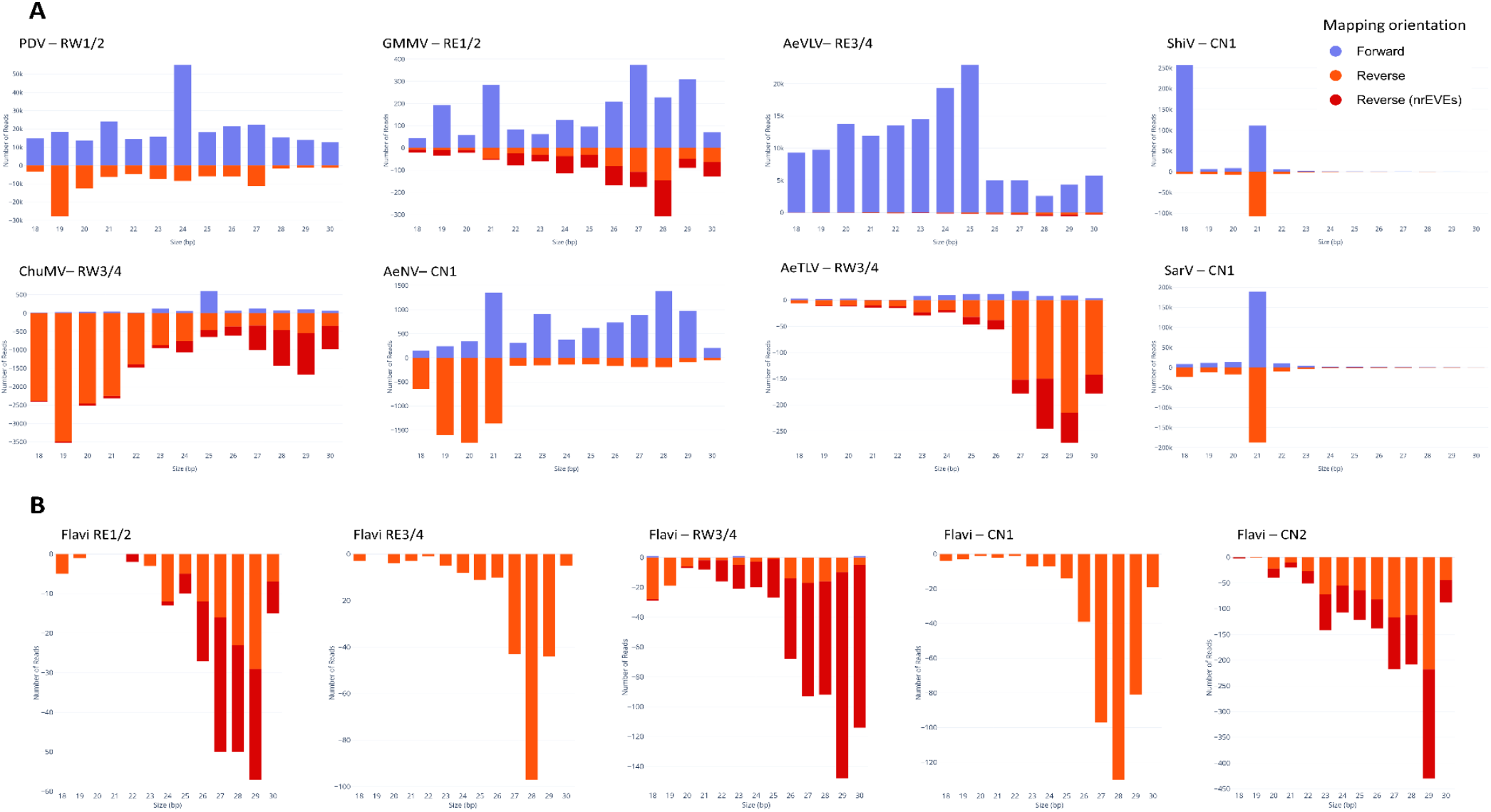
Profile of nucleotide lengths of small RNA reads mapping on viral contigs. (A) Profile of reads mapping on the 8 longest annotated viral contigs. (B) Profile of reads mapping to flaviviral contigs. piRNA -size read which also map to nrEVEs are shown in dark red. On top of each panel the viral contig along with the sample name are indicated. PDV=Point-Douro narnavirus; GMMV=Guadeloupe mosquito mononega-like virus; AeVLV=Aedes aegypti virga-like virus; ShiV=Shinobi tetravirus; ChuMV=Chuvirus Mos8Chu0; AeNV=Aedes albopictus negev-like virus; AeTLV=Aedes aegypti toti-like virus; SarV=Sarawak Virus.

As our analyses were being completed, independent groups of investigators used a metagenomic approach to describe the virome of mosquitoes collected between 2017-2018 from urban Guangzhou (60) and that of *Aedes* mosquitoes using published RNA-seq datasets (9, 11). A total of 42% of the viral contigs we identified had already been detected in world-wide *Ae. albopictus* samples (**Fig. 3**), further validating small RNA seq based virome identification and suggesting the existence of a core virome.

## DISCUSSION

Eukaryotic genomes have integrated viral sequences, most of which derive from retroviruses encoding for retrotranscriptase and integrases. Previous work conducted by us, and others, revealed the presence of nrEVEs with similarity to, mostly, flaviviruses and rhabdoviruses in *Ae. albopictus* and *Ae. aegypti*, but not in the genomes of *Anopheles* species and *Culex quinquefasciatus* (61). Additionally, we have shown that the pattern of nrEVEs varies across geographic populations, with new integrations being detectable in the genome of wild-collected mosquitoes (20, 23). Here we report the identification of 7 novel nrEVEs in wild-collected *Ae. albopictus* mosquitoes. On average, these nrEVEs have a higher sequence identity to corresponding viral genomes in comparison to reference nrEVEs, suggesting more recent integration events. Newly identified *Ae. albopictus* nrEVEs have similarities to a range of cISFs, including CFAV, AeFV and KRV, unlike the new nrEVEs identified in *Ae. aegypti* wild mosquitoes (23). Additionally, newly identified nrEVEs appear broadly distributed across mosquitoes from different locations, probably because of the quick and chaotic global invasion of *Ae. albopictus* (12). Of note CFAV-2 and KRV-1 were exclusive of mosquitoes from Italy and La Reunion Island, among the sites we tested, suggesting that these integrations occurred after *Ae. albopictus* established on La Reunion island, which is a primary source of mosquito introductions to Italy (12). Small RNA profiles of newly identified nrEVEs confirm the presence of low abundance piRNAs, and not siRNAs, which appear in hotspots that are conserved across samples similarly to what is observed for reference nrEVEs of both *Ae. aegypti* and *Ae. albopictus* (20, 23). Overall, these results indicate that, independently of the timing of the integration event, piRNAs will derive from nrEVEs, but not throughout the nrEVE sequence. Thus, endeavors aiming at using nrEVEs in strategies for vector control either as landmarks for effector integration or taking advantage of their inherent antiviral activity against cognate viruses might require experimental validation of their piRNA profile prior transgenic applications (62).

Given the abundance of nrEVEs in the genome of *Aedes* mosquitoes, we further asked whether the presence of newly identified nrEVEs correlates with on-going viral infections and analyzed the virome of the same mosquitoes in which we tested for novel nrEVEs. While our samples could be grouped similarly based both on their nrEVE landscape and their virome (**Fig. 2**), indicating that sampling location is the main factor influencing both virome and nrEVEome, we did not assemble viral contigs from the same viruses for which novel nrEVEs were identified in any sample (**Fig. 3, 4**). These results along with the number of newly identified nrEVEs with respect to the number of wild mosquitoes we analyzed show that integration events are rare. Whether the load of infecting viruses or other ecological and/or biological circumstances such as thermal stress, co-infections with bacteria or fungi and/or the presence of active transposable elements are needed for an integration event to occur requires further investigations.

Overall, our small-RNA-based virome analysis led to the identification of 43 viruses; 18 of these viral species have already been described in *Aedes* spp. mosquitoes, with Phasi Charoen-like phasivirus, Guadeloupe mosquito monomega-like virus, Aedes anphevirus, toti-like viruses and Chuvirus Mos8Chu0 being the most prevalent (9, 11, 60). Detailed analysis of the smallRNA profiles of the assembled viral contigs showed a profile with 21 bp peaks in sense and antisense orientations indicative of active replication only for few viruses, such as Sarawak virus, Shinobi tetravirus and the Aedes albopictus negev-like virus. Consequently, we cannot exclude that some of the assembled viral contigs are simply related to the diet or are associated with mosquito microbiota. For instance, we assembled contigs with similarities to *Virgaviridae*, which are plant viruses, and *Totiviridae* for which fungi and protozoans are natural hosts, in both mosquitoes from China and La Reunion Island. Sarawak virus was first identified in 2017 from *Ae. albopictus* males collected in 2013 on the Sarawak island of Borneo and its genome sequence has been produced showing it is an *Alphatetraviridae* with three open reading frames (ORFs) (63). Shinobi tetravirus was discovered in 2018 in *Ae. albopictus* C6/36 cells and its presence was shown to suppress replication of Zika virus(64). Shinobi tetravirus readily infects *Ae. albopictus* strains in laboratory conditions (65), but its prevalence in wild mosquitoes is uncertain. Aedes albopictus negev-like virus was discovered in 2020 in the *Ae. albopictus* Aa23 cell line, where Wolbachia wAlbB was shown to negatively affect its replication (66).

Overall, our results highlight the need not only to continue this type of studies to further understand the diversity of viruses infecting mosquitoes, but also to start introducing a functional characterization of these viruses and the effects of their infections on mosquitoes’ fitness and metabolism.

Distinct small RNA profiles were observed for additional viral contigs, including Chuvirus Mos8Chu0, which warrant further investigations. Chuviridae are a poorly characterized group of viruses, which have been recently identified through metagenomic analyses in several insects from different orders including Hemiptera, Diptera and Coleoptera (67, 68). Insects in which Chuviridae have been identified tend to also harbor Chuviridae-like nrEVEs (25, 67), thus these viruses may represent an ideal model to study the arm-race with insects resulting in integration events.

## FIGURES

**Fig. S1.**
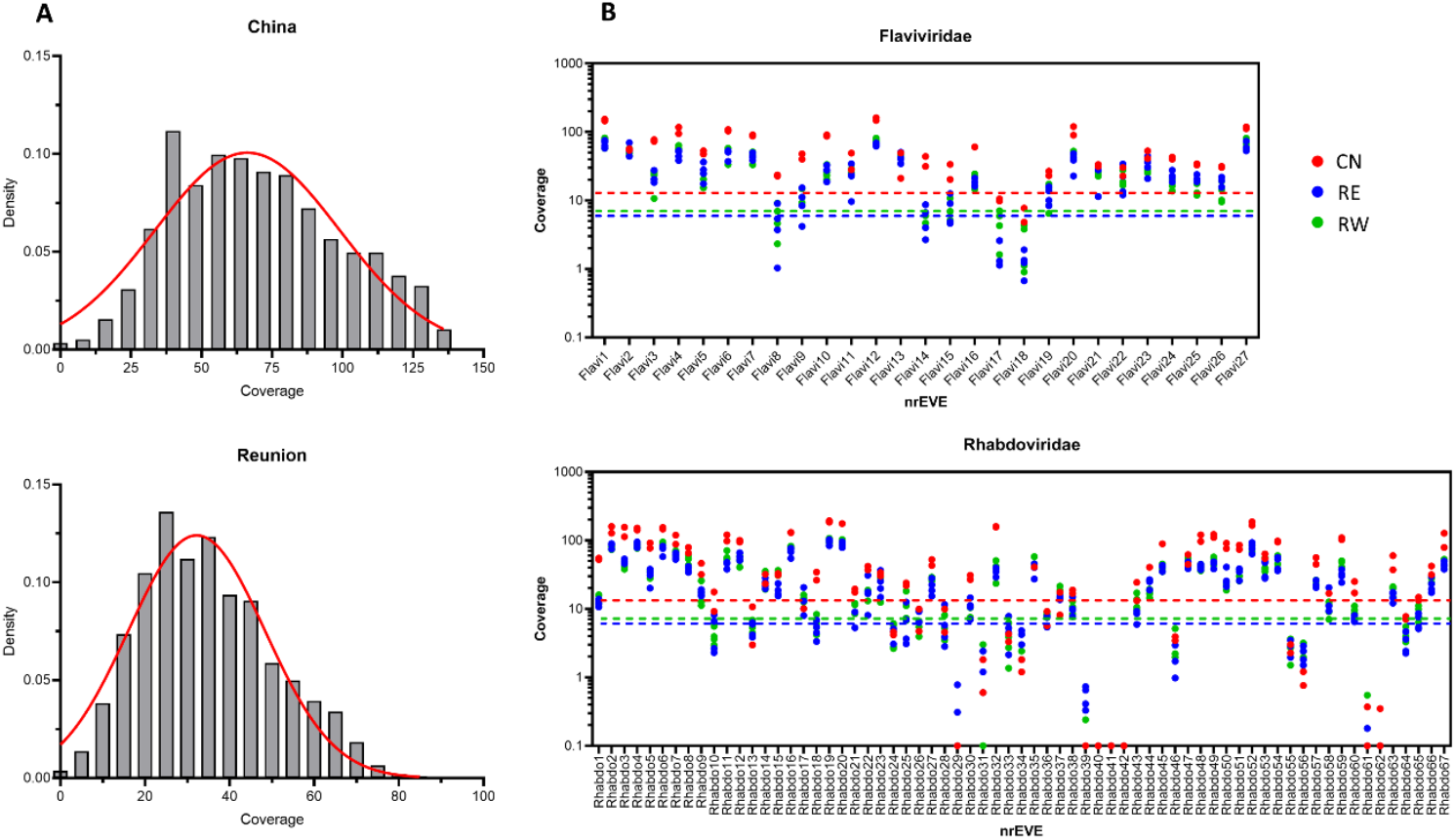
Presence or absence of AalbF2 annotated nrEVEs in samples from China and La Reunion. (A) Average distribution of reads coverage for 359 BUSCO genes calculated from the WGS mosquito data from China and La Reunion described in the study. (B) WGS reads coverage for *flaviviridaei*- and *rhabdoviridae*-derived annotated nrEVEs in samples from China (CN) and the Eastern and Western sides of La Reunion Island(RE and RW, respectively).

